# A humanized 16A antibody conjugated with DNA topoisomerase I inhibitors, targeting a GSTA glycosite-signature epitope

**DOI:** 10.1101/2025.03.22.644720

**Authors:** Yu He, Zhaohui Ji, Nian Wang, Jinxia Wang, Chenxi Zhu, Chunchao Feng, Xiaoding Tan, Dapeng Zhou

## Abstract

**Background:** We previously reported the 16A antibody, which binds to the abnormally glycosylated tandem repeat region of the MUC1 glycoprotein, and developed 16A-MMAE as an antibody-drug conjugate. However, its antitumor efficacy as an antibody-drug conjugate with DNA topoisomerase Iinhibitors remains unknown.

**Methods:** We humanized the 16A antibody and conjugated it to DXd and MF6, two DNA topoisomerase I inhibitors. The antitumor efficacy of the conjugates was evaluated in vitro and in vivo.

**Results:** The humanized 16A-DXd conjugate showed potent antitumoral efficacy, with an IC50 in the nM range against CFPAC1 cells. In vivo, h16A-DXd and h16A-MF6 inhibited tumor growth in asubcutaneous mouse tumor transplantation model using CT26-COSMC KO-hMUC1 cells, administered at a dose of 10 mg/kg.

**Conclusions:** The high antitumor efficacy of h16A-conjugated DNA topoisomerase I inhibitors supports further clinical development. The relatively low toxicity of h16A-DXd and h16A-MF6 may enable a higher therapeutic window, since MUC1-positive cells internalize the antibody-drug conjugate more efficiently.

## Introduction

MUC1 is a major glycoprotein carrying abnormally glycosylated glycans (1). It has been a target for the development of antibody-drug conjugates (ADCs) by major pharmaceutical companies (2-5). Both high-toxicity payloads like DM4 (used in SAR566658; *J Clin Oncol*. 2016, 34, suppl; abstr 2511) and low-toxicity payloads like DXd (used in DS3939; *J Clin Oncol*. 2024, 42, suppl; abstr TPS3165) have been tested in clinical trials. For SAR566658 (anti-CA6-SPDB-DM4), the dose of 190 mg/m^2^ every 3 weeks (q3w) was subsequently reduced to 150 mg/m^2^ q3w due to a high incidence of keratitis. No toxicity data were reported for DS3939 (PankoMab-DXd), which has entered clinical trials globally, including in Japan, the US, and China.

We previously screened antibodies specific to glycopeptide motifs of MUC1 and selected to design 16A-MMAE based on antibody internalization by tumor cells (1,6). In this study, we conjugated DNA topoisomerase I inhibitors, including DXd (7) and MF6 (8), to the humanized 16A antibody and examined their antitumor efficacy in vitro and in vivo.

## Materials and Methods

### Cell lines and reagents

Human tumor cell lines NCI-H838 (H838), A549, MCF-7, CFPAC1, and PANC1 (obtained from the American Type Culture Collection [ATCC]) were cultured at 37 °C with 5% CO2 in RPMI-1640 or DMEM media (Life Technologies) supplemented with 10% fetal bovine serum (FBS; Life Technologies). The CT26 mouse colon cancer cell line was genetically edited using CRISPR-Cas9 technology to inactivate the COSMC gene, which encodes a critical chaperone protein for Core 1 O-linked glycansynthesis (1). The CT26-COSMC KO cell line was stably transfected with a human MUC1 gene as described (2). CCFG-DXd was purchased from MedChemExpress (China). CCFG-MF6 was prepared as described previously. All chemical reagents and solvents were purchased from Sinopharm Chemical Reagent Co. (Shanghai, China) or Sigma-Aldrich and used without further purification.

### Humanization of 16A Antibody

Humanized antibodies were designed by creating hybrid sequences that fused selected regions of the parental antibody sequence with human framework sequences. Using a 3D model, these humanized sequences were analyzed via computer modeling to identify sequences most likely to retain antigen binding. We designed a humanized light chain (VL2) and five humanized heavy chain sequences (VH1–VH5) for the murine 16A antibody’s VL and VH genes.

### Flow Cytometry Staining of Cancer Cell Lines

Cell surface MUC1 expression was assessed by flow cytometry. Cells were washed with PBS containing 2% BSA and incubated with the 16A (1) or humanized 16A antibody for 30 minutes at 4 °C. After washing, cells were incubated with PE-labeled anti-mouse IgG or PE-labeled anti-human IgG (1 µg/mL) for 30 minutes at 4 °C. Cells were analyzed using a FACS Calibur (BD, NJ), and data were processed with FlowJo software (v7.6).

### Expression of h16A Antibody

DNA sequences encoding the h16A light and heavy chains were transfected into Chinese hamster ovarian (CHO) cells. CHO cells expressing h16A were cultured in a bioreactor for two weeks using Dynamis™ (Gibco) as the basal medium and Cell Boost™ 7a/7b (HyClone) as the feed medium. Supernatants were collected, and h16A was purified via Protein A affinity chromatography, anion-exchange chromatography, cation-exchange chromatography, and nanofiltration.

### Synthesis of h16A-DXd-DAR4 and h16A-DXd-DAR8

h16A antibody was treated with tris(2-carboxyethyl)phosphine hydrochloride (TCEP HCl; Sigma-Aldrich) to reduce inter-chain disulfide bonds. The antibody was diluted to 10 ± 3 g/L in 10 mM histidine buffer (pH 6.0 ± 0.3). A 10 ± 2 mg/mL aqueous TCEP solution was added at a molar ratio of ≥8:1 (TCEP:antibody). mc-GGFG-DXd (dissolved in dimethylacetamide) was mixed with the reduced antibody. The drug-to-antibody ratio (DAR) was controlled by adjusting TCEP:antibody and GGFG-DXd:antibody ratios. Post-conjugation, residual TCEP and dimethylacetamide were removed via chromatography.

### Synthesis of Peptide-Linker Coupled with MF6

For MF6 conjugation, TCEP HCl was added to h16A at a 10:1 molar ratio and incubated at 25 °C for 120 minutes. Peptide-linker-MF6 was introduced at a 4.8:1 molar ratio (MF6:antibody) and incubated at 25 °C for 60 minutes. The ADC was purified by hydrophobic interaction chromatography (HIC), followed by gentle mixing at 35 °C for 120 minutes for hydrolysis. The final product was buffer-exchanged to 10 mM phosphate buffer (pH 7.4) via ultrafiltration/diafiltration.

### Hydrophobic Interaction Chromatography (HIC)

ADC samples (1.0 mg/mL) were loaded onto a SHIMSEN Ankylo HIC-Ph column (4.6 × 100 mm). Mobile phases:

- Phase A: 0.025 M sodium phosphate, 0.02 M tetrabutylammonium bromide, 1.2 M ammonium sulfate (pH 7.4).
- Phase B: 0.025 M sodium phosphate, 0.02 M tetrabutylammonium bromide (pH 7.4).
- Phase C: 100% isopropyl alcohol (IPA).

ADCs were eluted in a gradient mode at 0.6 mL/min (A280 detection). DAR species were quantified via area normalization.

### Native Mass Spectrometry Analysis

ADCs were deglycosylated with PNGase F (1 μ □ [500 U] per 100 μg sample; RHINO BIO, China) at 37 °C for 16 hours. Separation used a Thermo Scientific™ MAbPac™ SEC-1 column (4 × 300 mm, 5 μm, 300 Å) equilibrated with 200 mM ammonium acetate (0.15 mL/min). Eluent was analyzed on an Agilent 6545XT Q-TOF mass spectrometer (ESI gas: 10 L/min, 325 °C; fragmentor: 380 V; scan range: 500–9000 m/z; acquisition rate: 1 spectrum/s).

### Cytotoxicity Assay

Human cancer cell lines (CFPAC-1, A549, PANC-1, H838, MCF-7) were cultured in appropriate media, harvested by trypsinization, and seeded into 96-well plates (3,000, 1,000, 5,000, 1,000, and 5,000 cells/well, respectively). ADCs were added at serial concentrations and incubated for 6 days. Cell viability was quantified using CellCounting-Lite 3D Luminescent Reagent (Vazyme, #DD1102-02). Luminescence was measured on a Molecular Devices SpectraMax M5, and IC50 values were calculated via 4-parameter fitting (SoftMax Pro v7.0.2 GxP).

### Efficacy of h16A-DXd and h16A-MF6 in Mouse Tumor Models

Animal studies were approved by Tongji University’s Institutional Animal Care Committee (Shanghai, China). Balb/c *nu/nu* mice (6-week-old females) were inoculated subcutaneously with 5 × 10□ CT26-COSMC KO-hMUC1 cells in 200 μL PBS. Tumor volume (V = ½ × a × b^2^) was measured starting at day 8. Mice were randomized into groups (n=5) when tumors reached 25–40 mm^3^ and treated with:

- h16A (10 mg/kg)
- h16A-DXd-DAR4 (10 mg/kg)
- h16A-MF6-DAR4 (10 mg/kg)
- Vehicle (PBS)

Two ADC doses were administered intravenously (48-hour interval). Statistical analysis used GraphPad Prism v5.

## Results

### Specificity and Affinity of h16A Antibody

To determine the specificity of humanized 16A, we stained CT26 cells and CT26-COSMC KO cells transfected with human MUC1. COSMC is the key chaperone protein for Core 1 O-linked β3-galactosyltransferase-1, the enzyme responsible for synthesizing Core 1 O-glycans. The knockout cells express exposed Tn antigen (GalNAc) and sialyl-Tn (sialyl-GalNAc) residues, which are characteristic of cancer-associated glycoforms. h16A antibody showed 10- to 100-fold higher bindingto COSMC KO cells across tested concentrations.

### Drug-Antibody Ratio (DAR) of h16A-DXd and h16A-MF6

h16A-DXd ADCs were prepared with two DAR values (h16A-DXd-DAR4 and h16A-DXd-DAR8), while h16A-MF6 ADC was prepared at DAR4. DAR values were determined via hydrophobic interaction chromatography (HIC). As shown in Figure 2, h16A-DXd-DAR4 exhibited DAR species distributions of 2.5% (DAR0), 21.3% (DAR2), 50.3% (DAR4), 22.5% (DAR6), and 2.2% (DAR8), with an average DAR of 3.96. h16A-DXd-DAR8 and h16A-MF6-DAR4 showed 100% homogeneity at their target DAR values. Native mass spectrometry confirmed these results (Supplemental Figure 1).

**Figure 1.**
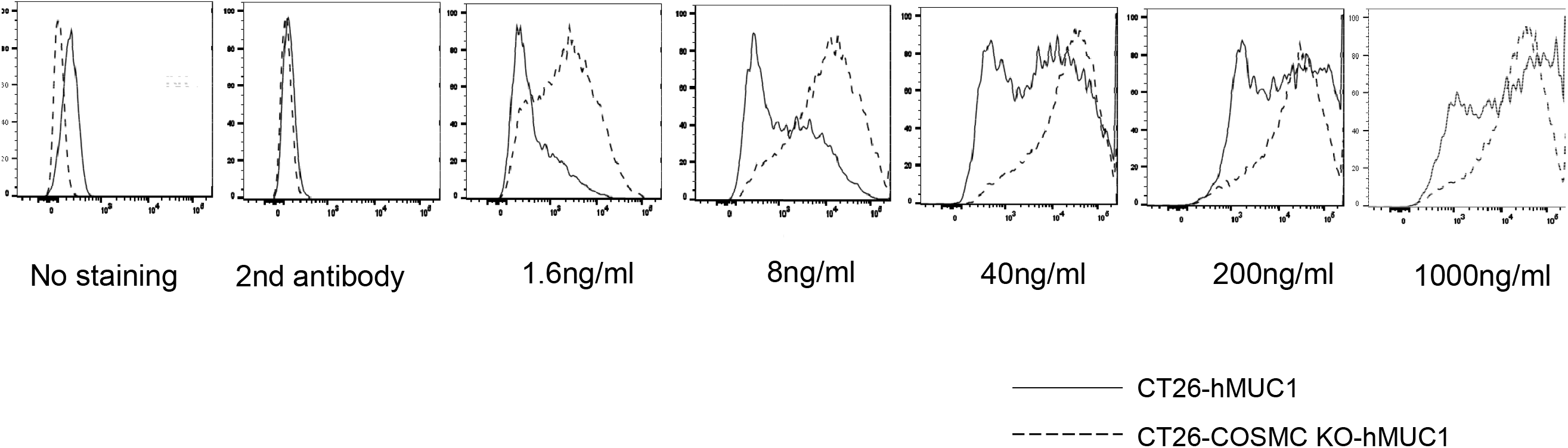
Specificity and affinity of h16A antibody. CT26 and CT26-COSMC KO cell lines were stably transfected with human MUC1 and stained with thehumanized 16A antibody.

**Figure 2.**
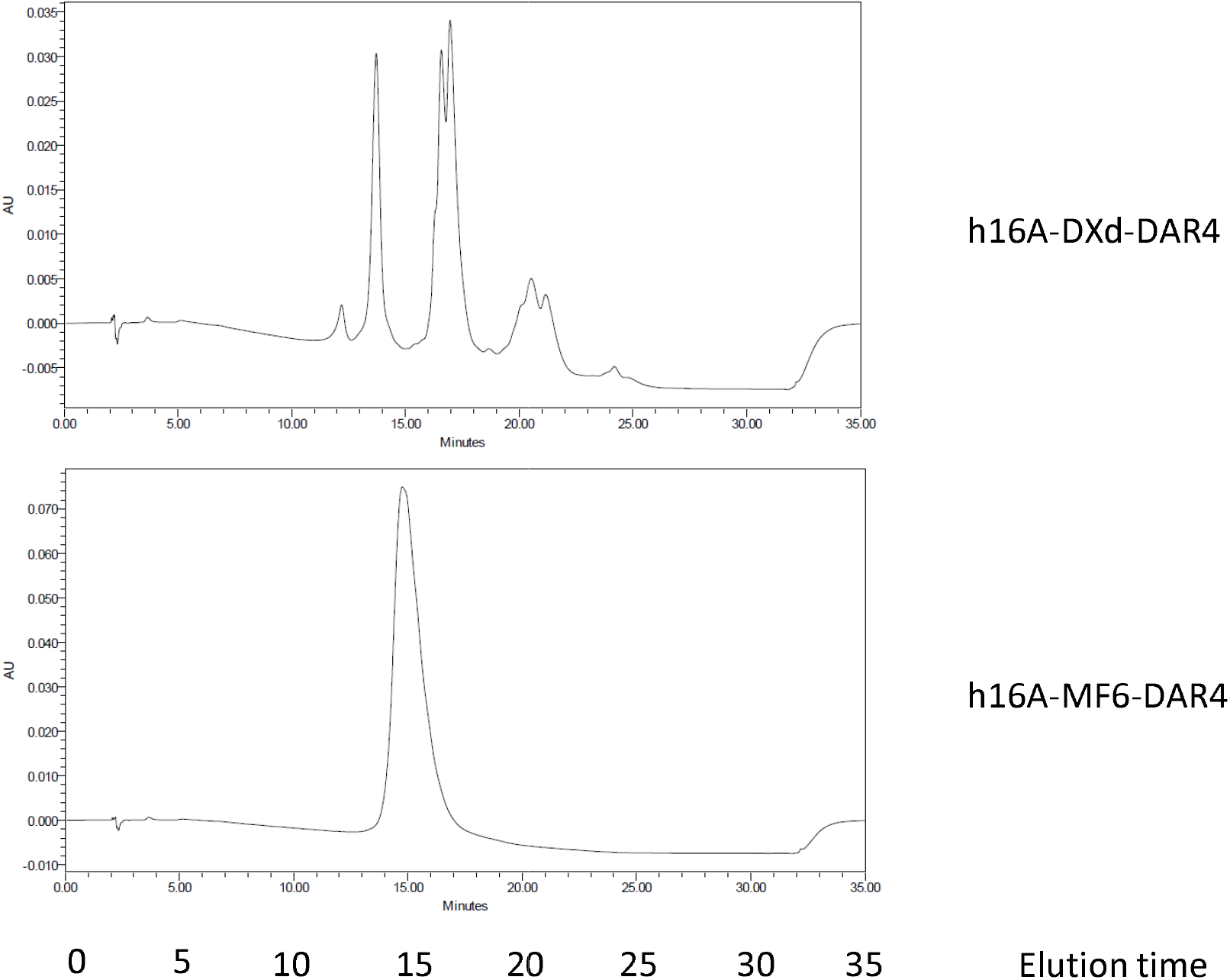
Drug-antibody ratio (DAR) of h16A-DXd-DAR4 and h16A-MF6-DAR4. The DAR of antibody-drug conjugates (ADCs) was measured by hydrophobic interaction chromatography (HIC).

### IC50 of h16A-DXd and h16A-MF6

The tumor-killing activity of h16A ADCs was assessed using cell viability assays. h16A-DXd demonstrated concentration- and DAR-dependent cytotoxicity against CFPAC-1, H838, and MCF-7 cells, but no significant cytotoxicity against A549 or PANC-1 cells. The IC50 values for h16A-DXd-DAR4 and h16A-DXd-DAR8 were 1.43 nM and 0.23 nM, respectively (Table 1).

**Table 1.** IC_50_ values of h16A-DXd for cancer cell lines.

| Cell lines | h16A-DXd-DAR4 (nM) | 16A-DXd-DAR8 (nM) |
| --- | --- | --- |
| A549 | No significant killing |  |
| H838 | Cell was killed without IC<sub>50</sub> due to curve fitting failure |  |
| MCF-7 | Cell was killed without IC<sub>50</sub> due to curve fitting failure |  |
| CFPAC1 | 1.43 | 0.23 |
| PANC-1 | No significant killing |  |

### In Vivo Antitumor Activity of h16A-DXd and h16A-MF6

The antitumor efficacy of h16A-DXd-DAR4 and h16A-MF6-DAR4 was evaluated in a subcutaneous CT26-COSMC KO-hMUC1 mouse tumor model. Both ADCs significantly inhibited tumor growth (Figure 3). Complete tumor eradication was achieved at a total dose of 10 mg/kg (administered as two 5 mg/kg doses).

**Figure 3.**
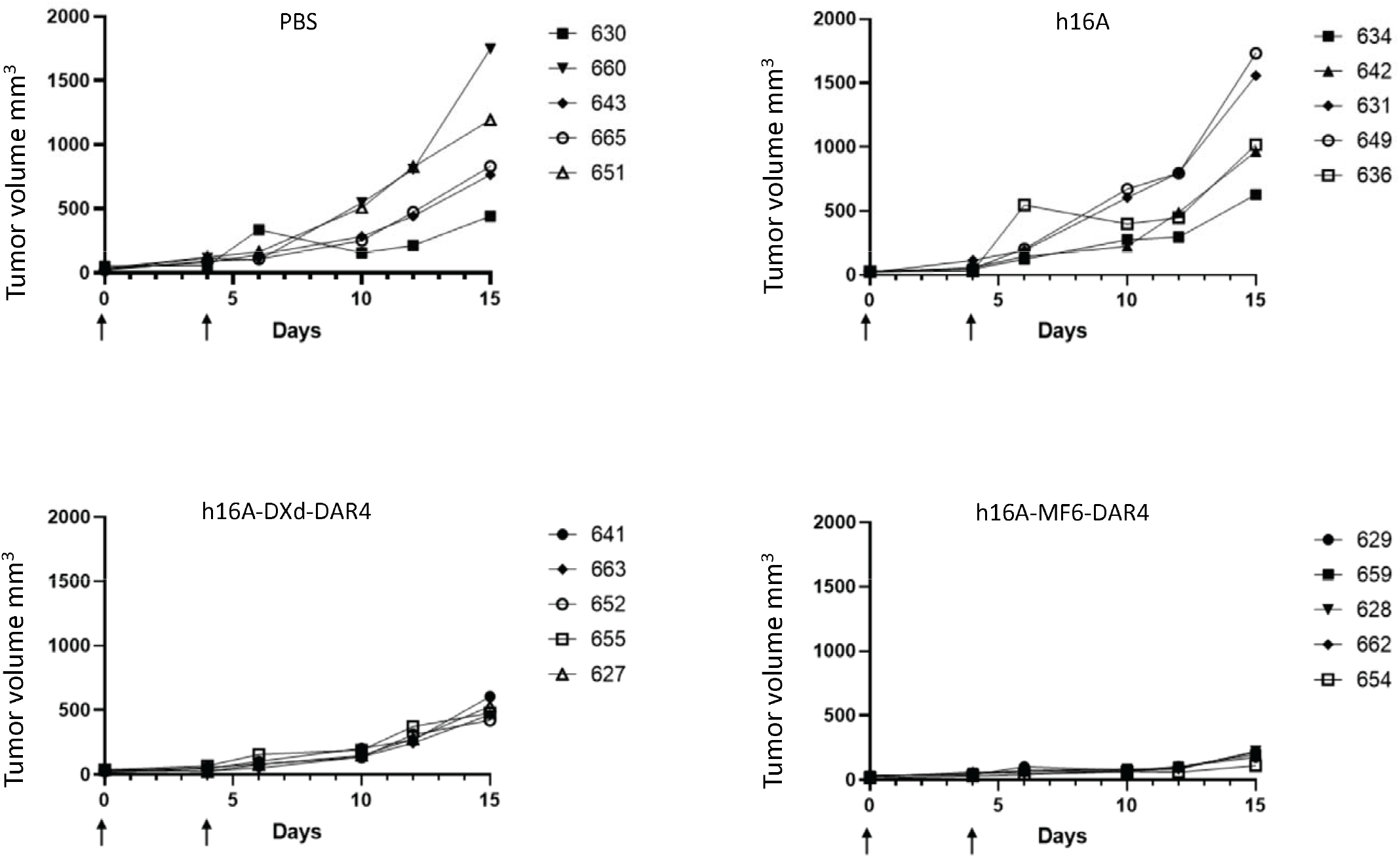
In vivo antitumor activity of h16A-DXd and h16A-MF6. The antitumor efficacy of h16A-DXd-DAR4 and h16A-MF6-DAR4 was evaluated in a subcutaneous mouse tumor model using CT26-COSMC KO-hMUC1 cells. Arrows indicate the days ADC was injected. Data was representative of three independent experiments.

## Discussion

### h16A-ADC’s Antitumor Efficacy and Cell Surface Expression of MUC1

Our previous studies demonstrated strong positivity of 16A staining in most cancer cell types. The number of MUC1 molecules expressed on tumor cells is highly variable, ranging from 10□ to 10□ across different cancer cell lines. Critically, the cell surface density of MUC1 determines the therapeutic efficacy of ADCs by modulating target engagement. We found that cancer cells with COSMC dysfunction (caused by mutations or hypermethylation of the COSMC gene [9–12]) bind humanized 16A antibody with 10- to 100-fold higher affinity (Figure 1). This selectivity positions h16A as a unique carrier for cytotoxic payloads, targeting tumor cells with aberrant glycoforms while sparing healthy tissues. In ongoing clinical studies, □ □Zr-labeled h16A shows in vivo internalization in cancer patients but no binding to normal mucosal membranes (data to be published elsewhere).

### Cell Lines and In Vivo Models for h16A-DXd

We used the CT26 mouse colon cancer model, a well-established system for evaluating ADCs conjugated with DNA topoisomerase inhibitors. h16A-DXd-DAR4 exhibited high antitumor activity at 10 mg/kg, a dose within the safe range for DXd-based ADCs. We further developed MF6, a novel DNA topoisomerase inhibitor with greater potency than DXd (8).

Consistent with its enhanced activity, h16A-MF6-DAR4 achieved complete tumor eradication at 10 mg/kg.

## Summary

This study reports the development of a humanized 16A antibody-drug conjugate (ADC) armed with DNA topoisomerase inhibitors (DXd and MF6). These ADCs demonstrate potent antitumor efficacyand a broader therapeutic window compared to older payloads like MMAE and DM4, underscoring their promise for clinical translation.

## Author contributions

Dapeng Zhou and Xiaoding Tan designed this study. Yu He, Zhaohui Ji, Nian Wang, Jinxia Wang, Chenxi Zhu, Chunchao Feng, Dapeng Zhou, Xiaoding Tan contributed to the collection, analysis and interpretation of data. Xiaoding Tan and Dapeng Zhou wrote the manuscript. All authors read and approved the final manuscript.

## Figure legends

**Supplemental Figure 1. Native mass spectrometry analysis of ADCs**

Native mass spectrometry analysis of h16A-DXd-DAR4 and h16A-DXd-DAR8.

