## Supplementary figures and images for "A humanized 16A antibody conjugated with DNA topoisomerase I inhibitors, targeting a GSTA glycosite-signature epitope"

### Supplemental Figure 1

h16A-DXd-DAR4

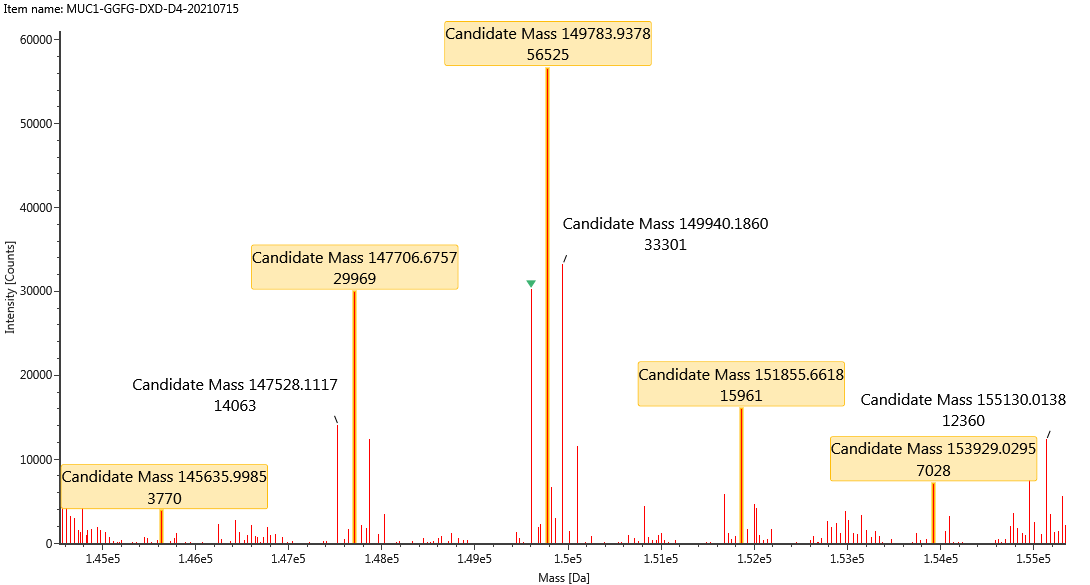
 h16A-DXd-DAR8
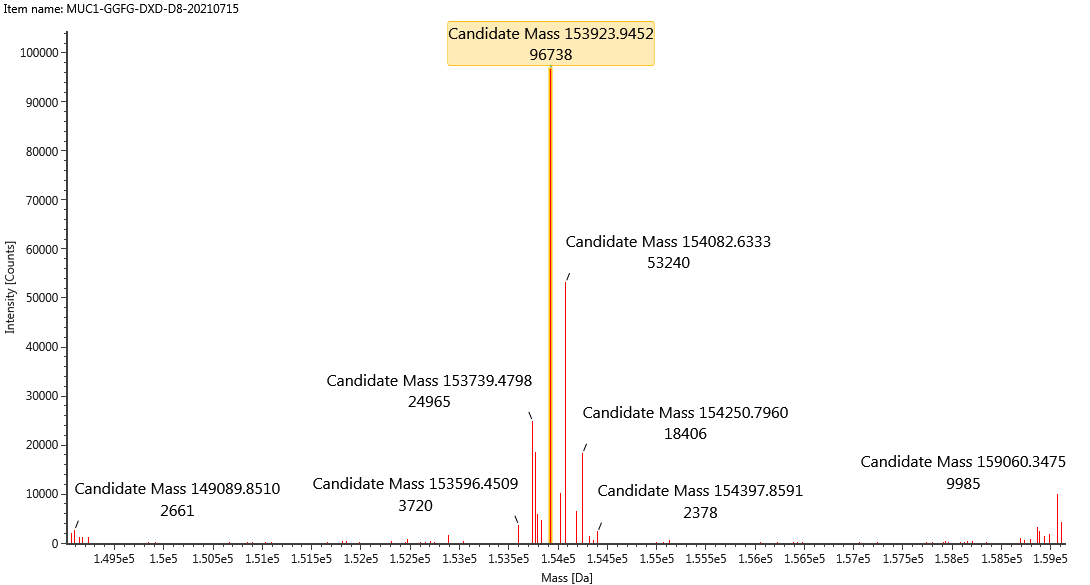
